# Augmented-reality-based multi-person exercise has more beneficial effects on mood state and oxytocin secretion than standard solitary exercise

**DOI:** 10.1101/2024.02.08.579578

**Authors:** Takeru Shima, Junpei Iijima, Hirotaka Sutoh, Chiho Terashima, Yuki Matsuura

## Abstract

Exercise is known to have positive effects on psychological well-being, with team sports often associated with superior mental health compared to individual sports. Augmented reality (AR) technology has the potential to convert solitary exercise into multi-person exercise. Given the role of oxytocin in mediating the psychological benefits of exercise and sports, this study aimed to investigate the impact of AR-based multi-person exercise on mood and salivary oxytocin levels. Fourteen participants underwent three distinct regimens: non-exercise (Rest), standard solitary cycling exercise (Ex), and AR-based multi-person cycling exercise (Ex+AR). In both exercise conditions (Ex and Ex+AR), participants engaged in cycling at a self-regulated pace to maintain a Rating of Perceived Exertion of 10. In the Ex+AR condition, participants’ avatars were projected onto a tablet screen, allowing them to cycle alongside ten other virtual avatars in an AR environment. Mood states (assessed using POMS2) and saliva samples were collected before and immediately after each 10-minute regimen. Subsequently, the levels of salivary oxytocin were measured. Participants exhibited higher cycling speeds during Ex+AR compared to Ex, despite comparable levels of self-reported fatigue between the two groups. Notably, only the Ex+AR condition significantly improved mood states associated with depression-dejection and exhibited a trend toward suppressing anger-hostility in participants. Moreover, the Ex+AR condition led to a significant elevation in salivary oxytocin levels, while the Ex condition showed a trend toward an increase. However, changes in salivary oxytocin did not show a significant correlation with changes in mood states. These findings suggest that Ex+AR enhances mood states and promotes oxytocin release. AR-based multi-person exercise may offer greater psychological benefits compared to standard solitary exercise, although the relationship between oxytocin and mood changes remains inconclusive.

**Highlights:**

- Exercise, especially team sports, has positive effects on mood state, and oxytocin partially supports these effects.
- Augmented reality (AR) technology can potentially convert standard solitary exercise into multi-person exercise.
- AR-based multi-person exercise (Ex+AR) significantly ameliorated depression-dejection and increased salivary oxytocin levels, but standard solitary exercise did not.
- Ex+AR may provide more significant psychological benefits than standard solitary exercise.

## 1. Introduction

Numerous studies have underscored the advantageous impacts of exercise on both physiological and psychological well-being. Consequently, exercise is viewed as a therapeutic intervention not only for metabolic disturbances but also for mental disorders (Deslandes et al., 2009; Hwang et al., 2023; Laird et al., 2023; Viana and de Lira, 2020). The prevalence of mental disorders has been on the rise in recent decades (ten Have et al., 2023), with children and young adults under 24 years of age being particularly affected by the adverse mental health effects of the COVID-19 pandemic (Kauhanen et al., 2023). While team sport athletes typically exhibit lower levels of anxiety and depression compared to individual sport athletes (Pluhar et al., 2019), the mental well-being of team sport athletes is susceptible to the impact of the COVID-19 pandemic due to the lack of social team activities (Salles et al., 2022). Consequently, exercise modalities resembling team sports that can be practiced under any environmental condition, such as during the COVID-19 pandemic, are deemed crucial for sustaining and improving human mental health.

Extended Reality (XR) technology holds promise for facilitating exercise under various environmental conditions. XR technology encompasses a spectrum of natural and virtual environments, including Virtual Reality (VR), Augmented Reality (AR), and Mixed Reality (MR) technologies. XR-based communication is likely effective in enhancing interaction (Dong et al., 2013; Rogers et al., 2022), and its applications to enhance human capabilities are burgeoning across various fields, such as sports and medical science (Barteit et al., 2021; Janssen et al., 2023; Le Noury et al., 2023). Among XR technologies, AR technology is particularly suited for sports applications as it can be utilized without the need for a head-mounted display. Leveraging AR has the potential to convert standard solitary exercise into multi-person exercise. Nevertheless, research investigating the effectiveness of AR technology for multi-person exercise is still in its nascent stages.

Oxytocin is implicated in the psychological benefits derived from exercise and sports. Oxytocin is a nine-amino acid peptide synthesized in the paraventricular nuclei and supraoptic nuclei of the hypothalamus, subsequently secreted from the posterior pituitary gland into circulation. Social behaviors such as affectionate touch and physical proximity are known to stimulate oxytocin secretion (Burenkova et al., 2023; Caicedo Mera et al., 2021; Holt-Lunstad et al., 2008; Landgraf and Neumann, 2004). Light-intensity acute exercise has been shown to enhance mood and prosocial skills in healthy individuals (Basso and Suzuki, 2017; Edwards et al., 2017; Ludyga et al., 2022), with oxytocin believed to partially mediate these effects (Ludyga et al., 2022). Moreover, even brief 10-minute exercise sessions have been shown to elevate salivary oxytocin levels (de Jong et al., 2015). Given that variations in oxytocin secretion are influenced by the type of exercise (Rassovsky et al., 2019), disparities in oxytocin secretion may contribute to the psychological benefits observed in team sports, such as reduced anxiety and depression levels compared to individual sports (Pluhar et al., 2019).

Here, to investigate the beneficial effects of multi-person exercises using AR technology, we examined the alterations in mood state and salivary oxytocin levels across three experimental conditions: 1. AR-based multi-person cycling exercise (Ex+AR), 2. standard solitary cycling exercise (Ex), and 3. Rest.

## 2. Methods and materials

### 2.1. Participants

Fourteen Japanese college students (8 men and 6 women, age: 20.5 ± 1.7), who had received instructions and provided consent, participated in the study. Informed consent was obtained from all individual participants prior to their involvement. The Utsunomiya University Ethical Review Board approved the current investigation for Medical Research Involving Human Subjects (ethical number: H23-0012). Participants had no history of neurological, psychiatric, or respiratory disorders, diabetes, anemia, or other medical conditions. Additionally, all participants underwent Ex+AR training prior to the main experiment. Post hoc sensitivity analysis conducted on the current sample, with 80% power and α = 0.05, revealed adequate sensitivity to detect repeated-measures effects exceeding *f* = 0.63 and paired t-test differences exceeding *d* = 0.81 (with a two-tailed α), as calculated using G*Power ver. 3.1.9.6 (The G*Power Team).

### 2.2. Experimental procedure

All participants were randomly assigned to undergo all three experimental conditions at one-week intervals: rest, Ex, and Ex+AR. Participants were instructed to refrain from exhaustive exercise, as well as from consuming alcohol and caffeine, for at least 24 hours prior to each experiment. The experimental procedure is shown in Figure 1. Paired participants were simultaneously exposed to the experiment under the same conditions in separate rooms. Participants completed the POMS2 questionnaire, and 1 ml saliva samples were collected after a 10-minute rest period. Following another 10-minute rest, participants engaged in a 10-minute sedentary period, Ex, or Ex+AR. Subsequently, participants completed the POMS2 questionnaire again, and saliva samples were collected. Heart rate (HR) was measured using an HR sensor (H10; Polar Electro, Japan).

**Fig. 1.**
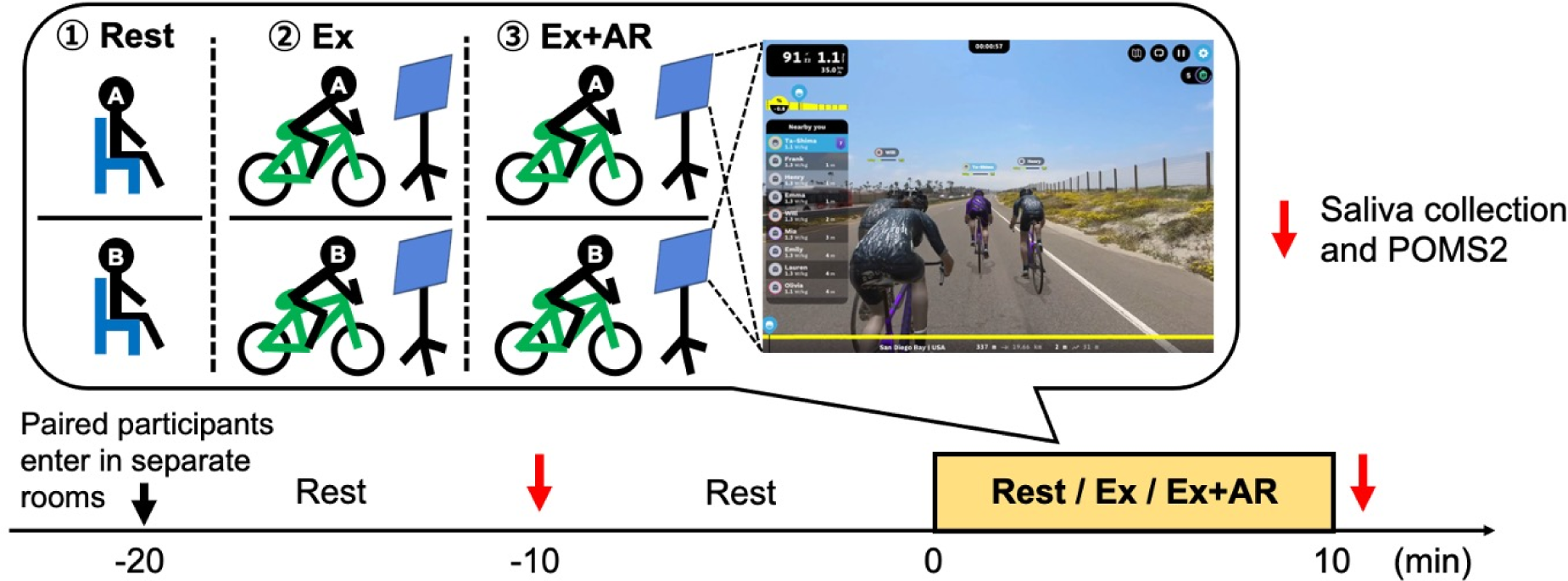
Experimental protocol in the current study.

### 2.3. Exercise and augmented reality system

In both exercise conditions (Ex and Ex+AR), participants engaged in self-paced cycling exercises to maintain a Rating of Perceived Exertion (RPE) of 10. Information regarding time remaining and confirming RPE remained at 10 were received, with no other information or external encouragement provided. Mean speed (km/h) was recorded throughout the 10-minute cycling session using the SmarT Turbo KAGURA (LST9200, MINOURA Co., Ltd., Japan) and the ROUVY application (VirtualTraining Co., Ltd., Czech Republic). During the Ex condition, participants cycled in front of a flipped tablet. In contrast, during the Ex+AR condition, participants cycled in front of a tablet (11-in) running the ROUVY application. In the Ex+AR condition, participants’ avatars were projected onto the tablet screen as they cycled in augmented reality alongside ten other avatars, including the avatar of a paired participant (refer to Fig. 1). Notably, participants did not utilize a headset with the current AR system.

### 2.4. Questionnaire

To assess the seven subscales of mood states (anger-hostility, confusion-bewilderment, depression-ejection, fatigue-inertia, tension-anxiety, vigor-activity, and friendliness), as well as total mood disturbance, participants completed the short form of POMS2 (Kaneko Shobo, Japan) before and immediately after each experimental condition. The short form of POMS2 comprises 35 items, with participants responding to a 5-point Likert scale ranging from 1 (strongly disagree) to 5 (strongly agree).

### 2.5. Saliva collection and measurement of oxytocin concentration

Saliva was chosen as the medium for evaluating oxytocin levels as it provides a reliable indicator of changes associated with interventions (Burenkova et al., 2023). Saliva samples were collected from participants before and immediately after engaging in each experimental condition using the Saliva Collection Aid (Salimetrics, LLC., USA) and Cryovial (Salimetrics, LLC., USA). These samples were stored at −20°C for subsequent analysis. Oxytocin concentrations in the saliva samples were measured in duplicate and calculated following the manufacturer’s instructions utilizing the Oxytocin ELISA Kit Wako (FUJIFILM Wako Pure Chemicals Corp., Japan) and a plate reader (EnSpire® Multimode Plate Reader, PerkinElmer, Inc., Massachusetts, USA).

### 2.6. Statistical analysis

The data are presented as mean ± SD and were analyzed using Prism version 10.1.1 (MDF, Japan). Before the analysis for group comparisons, we checked sphericity of the data. If the sphericity assumption was confirmed, group comparisons were conducted using repeated measures two-way ANOVA (factor 1: time points [pre vs. post]; factor 2: groups [rest vs. Ex vs. Ex+AR]) with Šídák’s multiple comparisons test. When the sphericity assumption was not confirmed, Geisser-Greenhouse correction was applied. A comparison between Ex and Ex+AR was performed using a paired t-test with checking the normality of raw data distribution using histograms (for analyzing mean speed). The correlation of mean speeds and changes in mood states with the changes in salivary oxytocin levels (baseline-normalized changes) were assessed using Pearson correlation. Changes in mood states were calculated by subtracting the scores at pre-exercise from the scores at post-exercise. Baseline-normalized changes in salivary oxytocin levels were calculated by subtracting the scores at pre from the scores at poste and then dividing by the scores at pre-exercise. Statistical significance was set at *p* < 0.05.

## 3. Results

### 3.1. Heart rate and mean speed during cycling exercise

Both the Ex and Ex+AR conditions demonstrated a significant increase in participants’ heart rate post-exercise compared to pre-exercise (Fig. 2A, Ex: *p* < 0.0001, Ex+AR: *p* < 0.0001; main effects of time points: *F*_(1.00, 13.00)_ = 129.0, *p* < 0.0001, main effects of group: *F*_(1.90, 24.70)_ = 40.6, *p* < 0.0001, interaction: *F*_(1.52, 19.71)_ = 79.8, *p* < 0.0001). Conversely, participants’ heart rate remained unchanged during the rest condition (Fig. 2A, *p* = 0.9984). Despite efforts to maintain an RPE of 10 in both the Ex and Ex+AR conditions, the mean cycling speed was higher in the Ex+AR condition compared to the Ex condition (Fig. 2B; *t* = 2.39, *p* = 0.033).

**Fig. 2.**
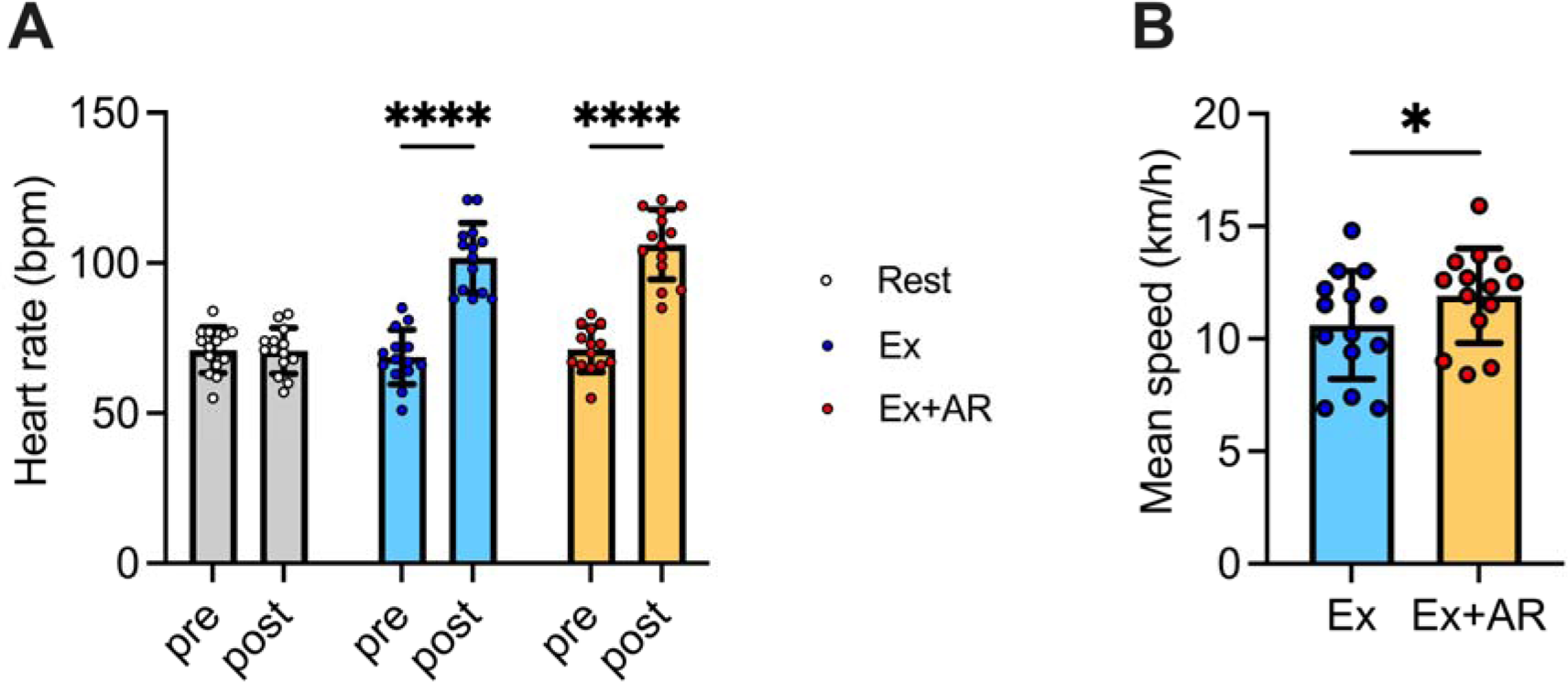
(A) The changes of heart rate in each condition. (B) The difference of mean cycling speed between Ex and Ex+AR condition. Ex: standard solitary cycling exercise, Ex+AR: augmented-reality-based multi-person cycling exercise. Data are expressed as mean ± SD. Plots represent individual data, n = 14. \**p* < 0.05, \*\*\*\**p* < 0.0001.

### 3.2. The changes in mood state

Only Ex+AR significantly suppressed the depression-ejection score (Fig. 3C, *p* = 0.0314; main effects of time points: *F*_(1.00, 13.00)_ = 6.92, *p* = 0.0208, main effects of group: *F*_(1.89, 24.57)_ = 1.53, *p* = 0.2370, interaction: *F*_(1.61, 20.92)_ = 0.72, *p* = 0.4686) and exhibited a trend towards a decrease in the scores of anger-hostility and total mood disturbance (Fig. 3A, H; anger-hostility: main effects of time points: *F*_(1.00, 13.00)_ = 8.87, *p* = 0.0107, main effects of group: *F*_(1.46, 18.95)_ = 0.04, *p* = 0.9240, interaction: *F*_(1.63, 21.20)_ = 1.26, *p* = 0.2960; total mood disturbance: main effects of time points: *F*_(1.00, 13.00)_ = 7.08, *p* = 0.0196, main effects of group: *F*_(1.77, 23.05)_ = 0.95, *p* = 0.3907, interaction: *F*_(1.70, 22.04)_ = 1.32, *p* = 0.2830). Both Ex and Ex+AR showed a trend towards a decrease in the tension-anxiety score (Fig. 3E; main effects of time points: *F*_(1.00, 13.00)_ = 15.0, *p* = 0.0019, main effects of group: *F*_(1.88, 24.41)_ = 1.01, *p* = 0.3747, interaction: *F*_(1.65, 21.46)_ = 1.27, *p* = 0.2941). None of the subscales of mood state or total mood disturbance showed alterations in the rest condition (Fig. 3A-H).

**Fig. 3.**
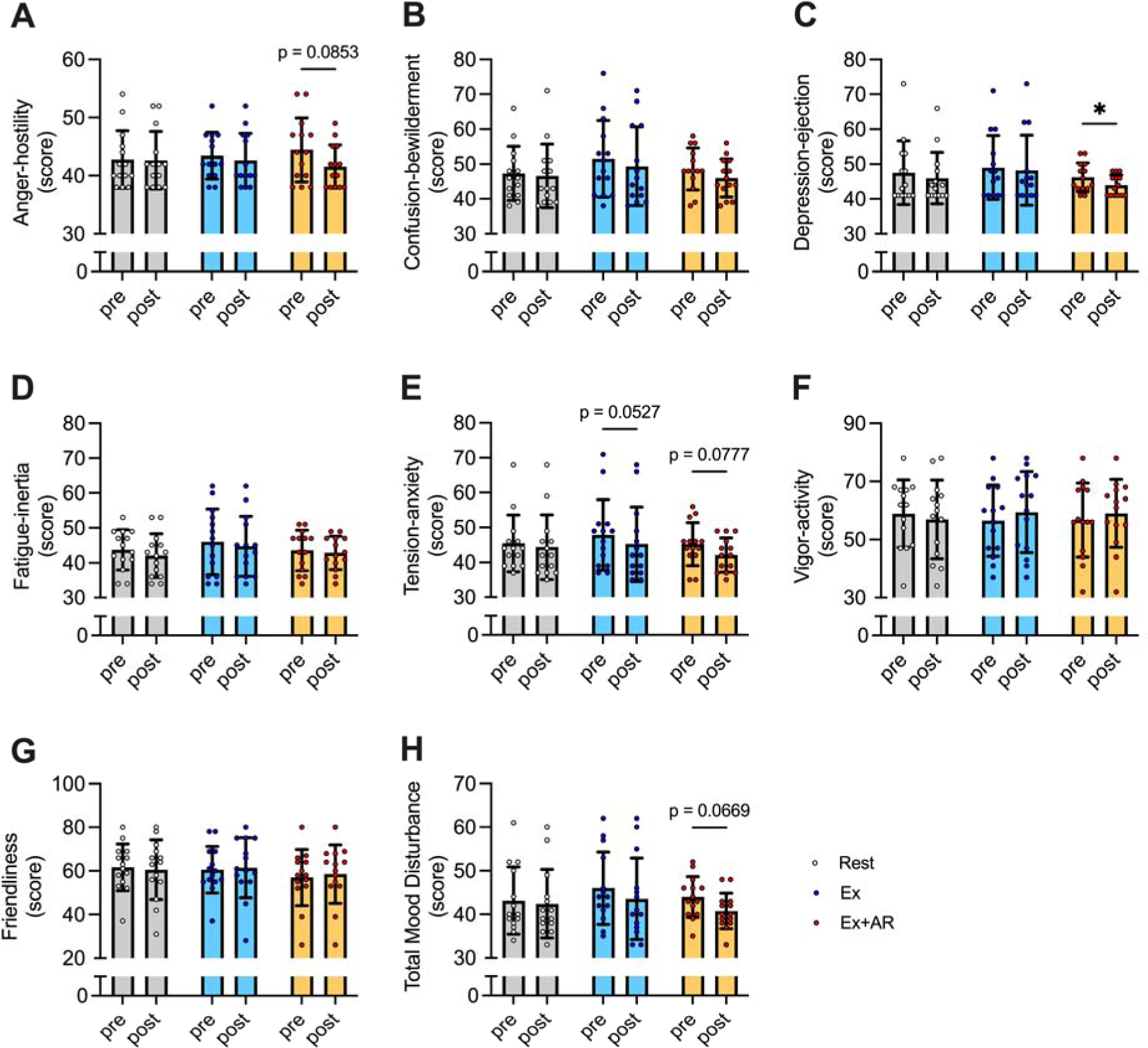
The changes of (A) anger-hostility, (B) confusion-bewilderment, (C) depression-ejection, (D) fatigue-inertia, (E) tension-anxiety, (F) vigor-activity, (G) friendliness and (H) total mood disturbance in each condition. Ex: standard solitary cycling exercise, Ex+AR: augmented-reality-based multi-person cycling exercise. Data are expressed as mean ± SD. Plots represent individual data, n = 14. \**p* < 0.05.

### 3.3. The changes in salivary oxytocin

Ex+AR significantly increased salivary oxytocin levels (Fig. 4A, *p* = 0.0005; main effects of time points: *F*_(1.00, 13.00)_ = 24.8, *p* = 0.0002, main effects of group: *F*_(1.84, 23.93)_ = 2.30, *p* = 0.1259, interaction: *F*_(1.94, 25.16)_ = 10.0, *p* = 0.0007), whereas Ex exhibited a trend towards an increase (Fig. 4A, *p* = 0.0868). Salivary oxytocin levels remained unchanged in the rest condition (Fig. 4A, *p* = 0.5238). On the other hand, no significant correlation was observed between cycling speeds and baseline-normalized changes in salivary oxytocin (Fig. 4B; *r* = 0.0411, *p* = 0.8354). Additionally, baseline-normalized changes in salivary oxytocin showed no significant correlation with changes in any mood states (Fig. 5A-H; all *p* > 0.05)

**Fig. 4.**
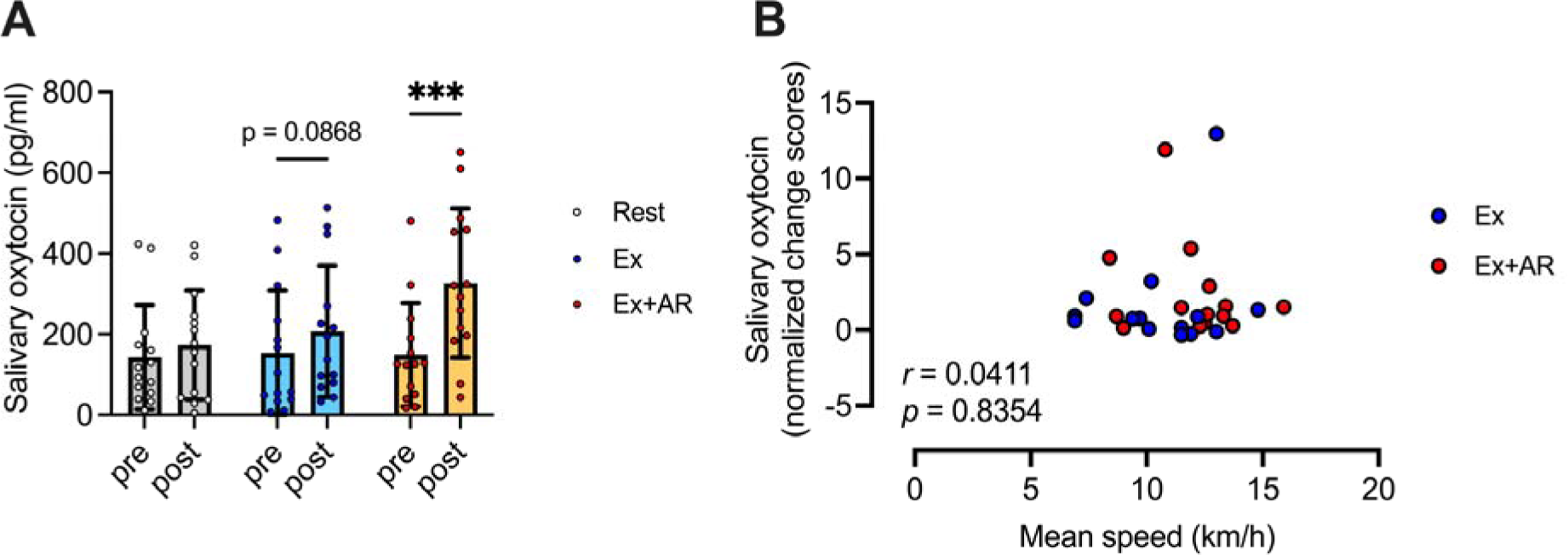
(A) The changes of salivary oxytocin levels in each condition. Ex: standard solitary cycling exercise, Ex+AR: augmented-reality-based multi-person cycling exercise. Data are expressed as mean ± SD. Plots represent individual data, n = 14. \*\*\**p* < 0.001. (B) The correlations between cycling speeds and baseline-normalized changes in salivary oxytocin levels. Blue circles, the Ex condition; red circles, the Ex+AR condition.

**Fig. 5.**
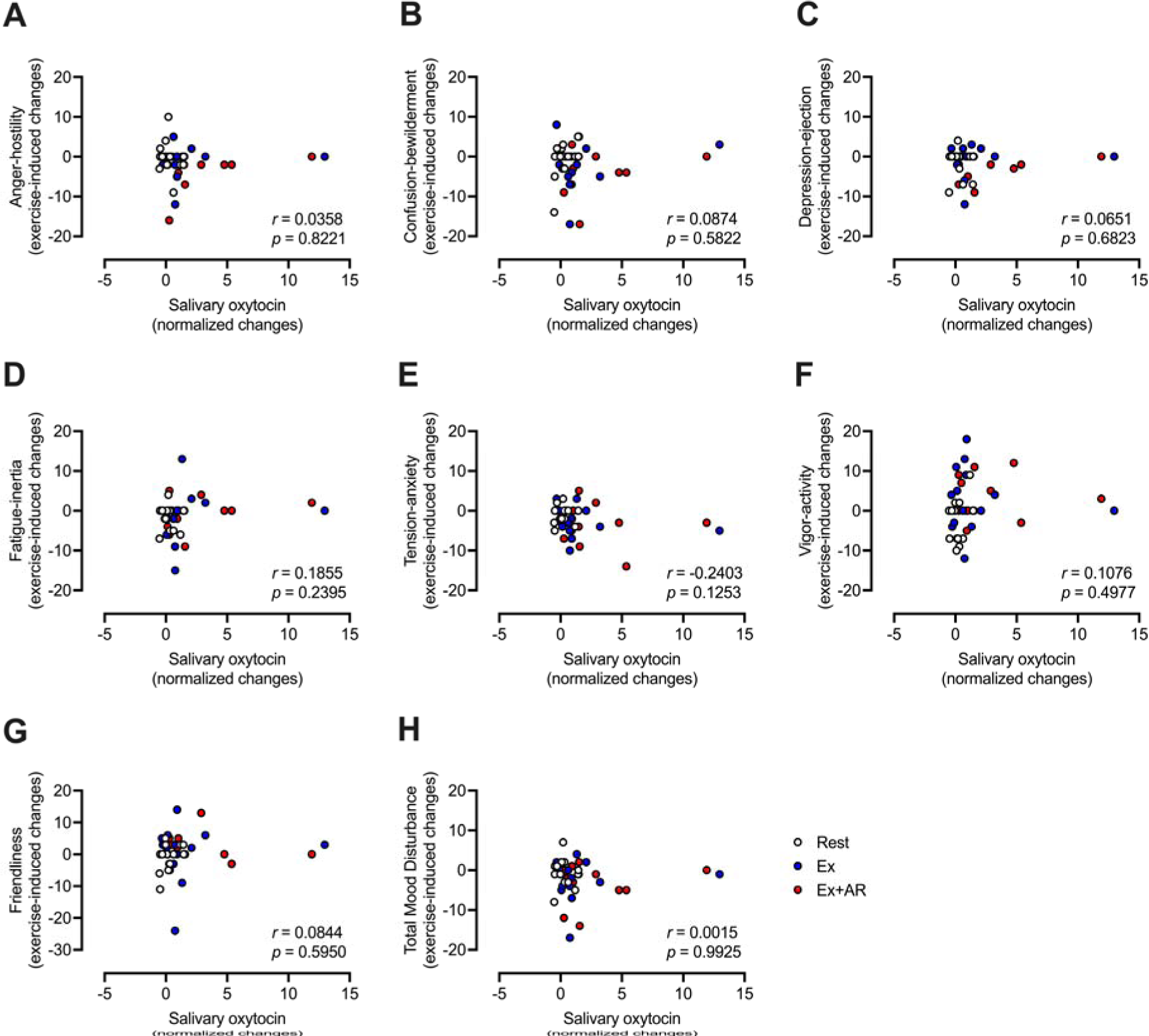
The correlations of baseline-normalized changes in salivary oxytocin levels with exercise-induced changes in (A) anger-hostility, (B) confusion-bewilderment, (C) depression-ejection, (D) fatigue-inertia, (E) tension-anxiety, (F) vigor-activity, (G) friendliness and (H) total mood disturbance in each condition. Ex: standard solitary cycling exercise, Ex+AR: augmented-reality-based multi-person cycling exercise. White circles, the Rest condition; blue circles, the Ex condition; red circles, the Ex+AR condition.

## 4. Discussion

In the current study, we investigated the effects of Ex+AR on psychological state and salivary oxytocin concentration compared to Ex. Our findings demonstrated that, despite comparable levels of effort to Ex, Ex+AR significantly improved depression-dejection scores and elevated salivary oxytocin levels.

Although participants engaged in cycling exercises at a self-regulated pace to maintain a RPE of 10 in the current study, the mean speed during cycling exercise in the Ex+AR condition was significantly faster with no difference in heart rate than in the Ex condition (Fig. 2A and B). Prior research has indicated that utilizing Extended Reality during exercise can mitigate human fatigue (Berezina et al., 2022; Plante et al., 2003). The current results showed no difference in fatigue evaluated by POMS2 among the groups (Fig. 3D). Therefore, utilizing Augmented Reality during exercise could potentially increase exercise intensity while maintaining similar levels of fatigue as standard exercise. It’s also worth considering that using an AR system without a head-mounted display may affect perceptual functions differently (Livingston et al., 2013). Since our study did not incorporate a head-mounted display in the AR system, future research should examine the comparative effects of the presence or absence of such a head-mounted display on the effectiveness of AR-based exercises.

Only the Ex+AR condition significantly suppressed the depression-ejection score (Fig. 3C) and showed a trend towards reducing anger-hostility and total mood disturbance scores (Fig. 3A, H). Considering that team sport athletes often exhibit a lower risk of anxiety and depression compared to individual sport athletes (Pluhar et al., 2019), the online exercise with multiple participants (ghost riders) in our Ex+AR condition may have induced effects similar to those observed in team sports. On the other hand, both the Ex and Ex+AR conditions showed a trend towards decreasing tension-anxiety scores (Fig. 3E), indicating the effectiveness of not only Ex+AR but also Ex, consistent with previous studies (Basso and Suzuki, 2017; Edwards et al., 2017). based on our findings, Ex+AR appears to have more beneficial effects on psychological aspects than Ex. Therefore, AR-based multi-person exercise may contribute to maintaining better mental health compared to standard solitary exercise.

Salivary oxytocin levels increased significantly only in the Ex+AR condition, while the Ex condition exhibited a trend towards an increase (Fig. 4A). Since baseline-normalized changes in salivary oxytocin levels did not correlate with cycling speeds (Fig. 4B), it suggests that the use of the AR system could enhance oxytocin secretion during exercise. Previous research has demonstrated that XR-based communication facilitates social interaction (Dong et al., 2013; Rogers et al., 2022) and oxytocin secretion (Dekker et al., 2021). Although we could not investigate enhancements in the social interaction of subjects by the Ex+AR condition, AR-based multi-person exercise would enhance not only mood states but also salivary oxytocin levels.

Baseline-normalized changes in salivary oxytocin levels did not correlate with changes in any mood states (Fig. 5A-H). While oxytocin has been implicated in reducing depressed mood, anger expression, and anxiety state (Domes et al., 2007; Galbally et al., 2021; Goodin et al., 2015; Heinrichs et al., 2003; Yoon and Kim, 2022), our findings suggest that changes in oxytocin secretion with exercise may not directly relate to exercise-induced changes in mood states. Oxytocin is also associated with empathy (Meyer-Lindenberg et al., 2011), and several studies have reported that oxytocin administration enhances empathy and related behaviors in humans (Auyeung et al., 2015; Geng et al., 2018). Considering that physical activities also contribute to enhancing empathy (Shima et al., 2022a, 2022b, 2021c, 2021a, 2021b), it is conceivable that exercise-induced oxytocin secretion plays a role in enhancing empathy. Therefore, AR-based multi-person exercise may have more beneficial effects on enhancing empathy than standard exercise, warranting further investigation.

The current study has several limitations. Firstly, the current study exclusively involved healthy participants. Given that standard exercise has been shown to prevent and treat psychiatric disorders (Deslandes et al., 2009; Hwang et al., 2023; Laird et al., 2023; Viana and de Lira, 2020), there is a possibility that the current Ex+AR approach could serve as a more efficient therapy for such disorders compared to standard exercise alone. Future studies should explore this possibility Secondly, our investigation focused solely on the acute effects of a 10-minute regimen. Prior research suggests that prolonged exposure to AR (>20 min) is associated with cybersickness (Hughes et al., 2020). Therefore, further study is warranted to determine the optimal duration for AR-based multi-person exercise. Thirdly, the effects of oxytocin and exercise may vary among individuals (Shima et al., 2022a; Sikich et al., 2021). Therefore, further research is needed to determine whether the current Ex+AR condition effectively enhances human well-being. Lastly, the assessment of social interaction provided by the AR system was not evaluated. Future studies would benefit from analyzing the quality of social interaction using established methods (Lopes et al., 2004; Reis and Wheeler, 1991).

In conclusion, our findings exhibit that Ex+AR significantly enhances mood states and increases salivary oxytocin levels. AR-based multi-person exercise may indeed provide greater psychological benefits compared to standard solitary exercise, although the relationship between oxytocin and mood changes remains inconclusive.

## Data availability

The datasets in the current study are available from the corresponding author on reasonable request.

## Author contribution

TS: Conceptualization, Methodology, Resources, Investigation, Writing–original draft, Writing–review & editing, Funding acquisition; JI: Investigation, Writing–review & editing, HS: Investigation, Writing–review & editing, CT: Investigation, Writing–review & editing, YM: Conceptualization, Methodology, Writing–review & editing.

